# Biochemical characterization of mRNA capping enzyme from Faustovirus

**DOI:** 10.1101/2023.06.05.542502

**Authors:** S. Hong Chan, Christa N. Molé, Dillon Nye, Lili Mitchell, Nan Dai, Jackson Buss, Daniel W. Kneller, Joseph M. Whipple, G. Brett Robb

**Affiliations:** New England Biolabs, Inc. 240 County Road, Ipswich, MA 01938

## Abstract

The mammalian mRNA 5’ cap structures play important roles in cellular processes such as nuclear export, efficient translation, and evading cellular innate immune surveillance and regulating 5’- mediated mRNA turnover. Hence, installation of the proper 5’ cap is crucial in therapeutic applications of synthetic mRNA. The core 5’ cap structure, Cap-0, is generated by three sequential enzymatic activities: RNA 5’ triphosphatase, RNA guanylyltransferase, and cap N7-guanine methyltransferase. Vaccinia virus RNA capping enzyme (VCE) is a heterodimeric enzyme that has been widely used in synthetic mRNA research and manufacturing. The large subunit of VCE D1R exhibits a modular structure where each of the three structural domains possesses one of the three enzyme activities, whereas the small subunit D12L is required to activate the N7-guanine methyltransferase activity. Here we report the characterization of a single-subunit RNA capping enzyme from an amoeba giant virus. Faustovirus RNA capping enzyme (FCE) exhibits a modular array of catalytic domains in common with VCE and is highly efficient in generating the Cap-0 structure without an activation subunit. Phylogenetic analysis suggests that FCE and VCE are descended from a common ancestral capping enzyme. We found that compared to VCE, FCE exhibits higher specific activity, higher activity towards RNA containing secondary structures, and broader temperature range, properties favorable for synthetic mRNA manufacturing workflows.

## Introduction

The rapid and global deployment of synthetic mRNA vaccines against SARS-CoV-2 was the result of decades of research on several fronts (Dolgin 2021). The core technologies of synthetic mRNA are *in vitro* transcription (IVT) using T7 RNA polymerase (Davanloo et al. 1984) and 5’ cap incorporation using Vaccinia virus RNA capping enzyme (Martin and Moss 1976; Shuman and Moss 1990; Fuchs et al. 2016) or cap analogs (Henderson et al. 2021; Ishikawa et al. 2009). The 5’ cap structures perform important biological functions such as efficient translation of the mRNA, evasion of the innate immune surveillance systems, and 5’-mediated mRNA turnover (Ramanathan et al. 2016). Therefore, 5’ cap incorporation is considered one of the critical quality attributes (CQAs) in mRNA manufacturing (Daniel et al. 2022). Methods to assess the extent of 5’ cap incorporation have recently been published (Chan et al. 2022; Vlatkovic et al. 2022; Cairns et al. 2003; Beverly et al. 2016).

The mRNA 5’ cap structures were first identified by multiple research groups in the 1970’s (Wei et al. 1975; Moyer et al. 1975; Furuichi et al. 1975; Desrosiers et al. 1975; Furuichi 2015). In lower eukaryotes such as fungi, mRNA contains a 5’ cap structure (Cap-0) where a N7-methyl-guanosine is linked to the 5’ end of the RNA via a 5’-to-5’ triphosphate linkage (Figure 1A). In metazoans, the Cap-0 structure is further modified to form the Cap-1 structure where a methyl group is added to the 2’-O group of the first nucleotide of the RNA (Barbosa and Moss 1978). The Cap-1 structure has been shown to be an important factor in evading the innate immune surveillance systems in humans (Hyde and Diamond 2015; Goubau et al. 2014). The Cap-1 structure can be further modified to a Cap-2 structure, where the 2’-O groups of the first and second nucleotides of the RNA are methylated (Werner et al. 2011; Drazkowska et al. 2022; Smietanski et al. 2014) (Figure 1A). The Cap-2 structure has been implicated in further evasion of innate immune response over the lifetime of mRNA molecules (Despic and Jaffrey 2023).

**Figure 1.**
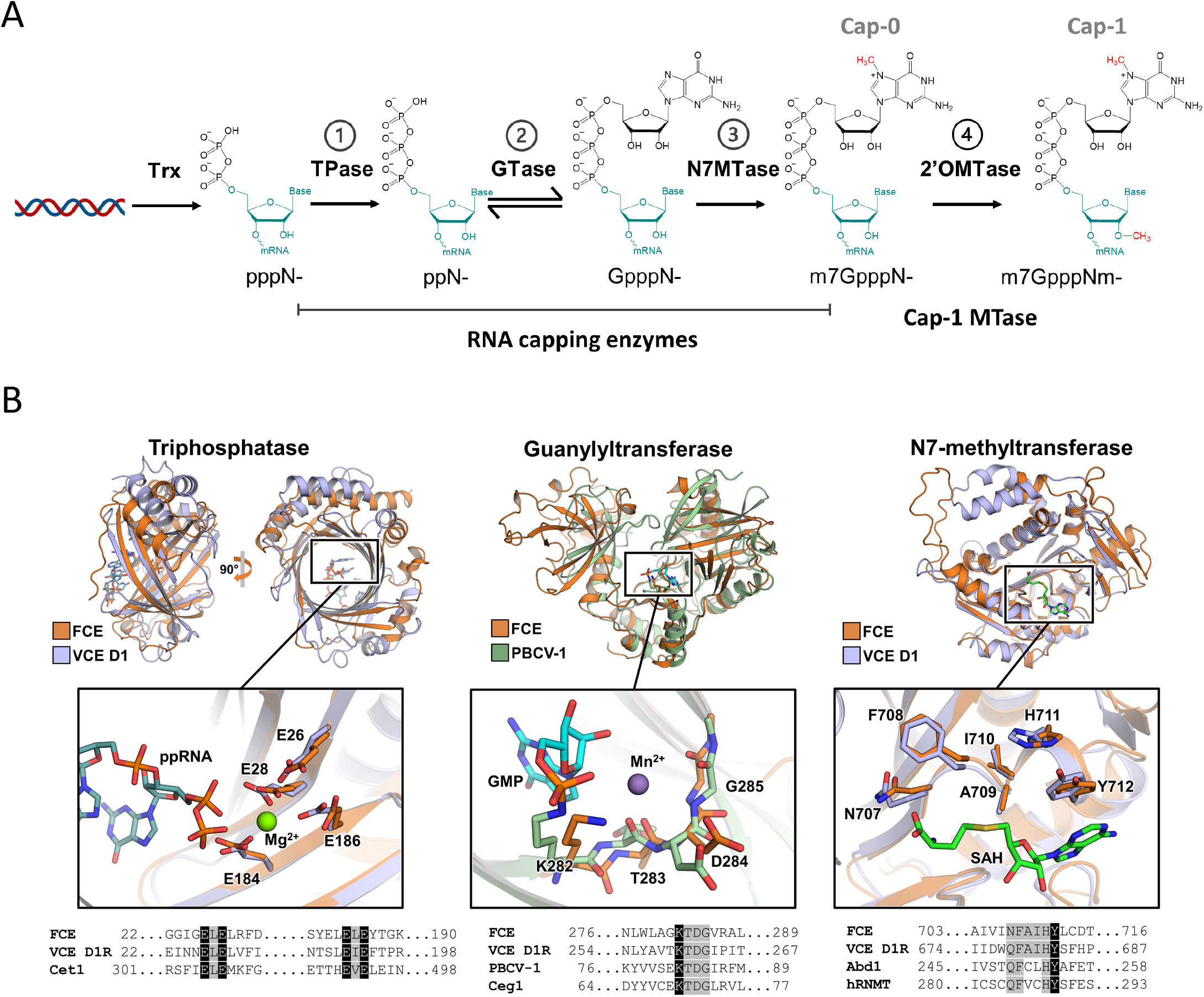
**(A)** Enzymatic reactions involved in RNA 5’ capping. First, the RNA triphosphatase (TPase) hydrolyzes the 5’ triphosphate group (pppN-) to form a 5’ diphosphate (ppN-) (Reaction 1). Then an RNA 5’ guanylyltransferase adds a GMP group to the 5’ diphosphate to form an unmethylated G-capped structure (GpppN-) through a reversible mechanism (Reaction 2). This is followed by an RNA cap guanosine N7-methyltransferase activity that adds a methyl group to the N7 position of the G cap and form the Cap-0 structure (m7GpppN-) (Reaction 3). In metazoans, a fourth enzyme RNA cap 2’-O-methyltransferase further adds a methyl group to the 2’-O position of the initiating nucleotide of the transcript, forming the Cap-1 structure (m7GpppNm-) (Reaction 4). **(B)** Catalytic domains of FCE. Top panels: Individual domains of AlphaFold2 prediction of FCE were aligned to published atomic structures. Insets show the alignment of highly conserved residues labeled with FCE residue numbering. Bottom panels: Amino acids residues of the motifs were aligned with indicated enzymes. Conserved residues are highlighted in grey. FCE residues that are mutated in this study and the corresponding residues in orthologous enzymes are highlighted in black. **Left column:** The predicted structure of FCE TPase domain is aligned to the VCE TPase domain-ppRNA complex extracted from the CryoEM co-transcriptional capping complex (PDB ID 6RIE). The inset shows the alignment of the conserved ELE-ELE motif. **Center column:** The predicted structure of FCE GTase domain is aligned to X-ray crystal structure of Paramecium bursaria chlorella virus (PBCV-1) GTase with covalent GMP intermediate (PDB ID 1CKN). Inset shows the phosphamide covalent linkage between the catalytic Lys of PBCV-1 GTase and GMP. **Right column:** The predicted FCE N7MTase domain structure is aligned to the X-ray crystal structure of the N7MTase domain of VCE D1 subunit in complex with co-product SAH (PDB ID 4CKB). Inset shows the conserved SAM binding site. Cet1, Ceg1 and Abd1 are the TPase, GTase and N7MTase of S. cerevisiae, respectively. hRNMT is the human RNA cap N7MTase.

The first mRNA capping enzyme was purified from Vaccinia virus particles (Martin and Moss 1975; Martin et al. 1975; Shuman et al. 1980) around the same time mRNA cap structures were discovered. As the first mRNA capping enzyme available in recombinant form (Shuman 1990; Shuman and Moss 1990), Vaccinia capping enzyme (VCE) became the reference for subsequent research on cellular and viral RNA capping enzymes (Ghosh and Lima 2010; Shuman 2015; De Vlugt et al. 2018; Park et al. 2022). For example, RNA capping enzymes in general refer to proteins that generate the Cap-0 structure through three enzyme activities that VCE possesses: the RNA triphosphatase activity (TPase), the RNA guanylyltransferase activity (GTase) and the RNA cap N7-guanosine methyltransferase activity (N7MTase) (Figure 1A, reactions 1-3). The TPase domain consists of the triphosphate tunnel metalloenzyme (TTM) fold where the 5’ triphosphate RNA is inserted into a long beta-barrel structure for hydrolysis (Kyrieleis et al. 2014). The GTase domain shares the adenosyl transferase active site of DNA and RNA ligases that involves a covalent Lys-GMP intermediate and has the propensity to catalyze the forward and reverse reactions, whereas the N7MTase has the characteristic core fold of class I MTase family (De la Peña et al. 2007; Kyrieleis et al. 2014).

The large subunit of VCE (D1R) exhibits a modular structure where each of the three structure domains carries one of the component enzyme activities (Kyrieleis et al. 2014; Shuman and Morham 1990). These functional domains are largely conserved in eukaryotes, but their organization can be variable. For example, three individual proteins, Cet1, Ceg1, and Abd1, carry each of the three activities in *S. cerevisiae* (Lima et al. 1999; Shuman et al. 1994; Mao et al. 1995). In mammals, RNMT possesses the N7MTase activity and RNGTT possesses the TPase and GTase activities where the TTM domain is replaced by a tyrosine phosphatase domain (Wen et al. 1998). In metazoans, the Cap-0 structure is further modified to the Cap-1 structure by a 2’-*O*- methyltransferase activity (2’OMTase) (Figure 1A, reaction 4). The enzymes CMTR1 and VP39 possess the 2’OMTase activity in humans and Vaccinia virus, respectively. RNA viruses have evolved a diverse set of mechanisms for mRNA capping: formation of an enzyme-pRNA covalent intermediate in vesicular stomatitis virus (Ogino and Banerjee 2007) and coronavirus (Park et al. 2022) through the kinase-like Nidovirus RdRp-associated nucleotidyltransferase domain (NiRAN); direct transfer of m7GMP to pp-RNA in Alphavirus (Ahola and Kääriäinen 1995; Tomar et al. 2011), and cap-snatching from host mRNA in Influenza virus and the yeast Totivirus (Ramanathan et al. 2016). Viral RNA capping mechanisms have been exploited for antiviral drug development. For example, cap-snatching inhibitors are an effective class of anti-influenza drugs (Todd et al. 2021; Hsu et al. 2012; García-Sastre 2019; Jones et al. 2016).

Vaccinia virus belongs to a monophyletic class of viruses called the nucleocytoplasmic large DNA viruses (NCLDV), most of which encode their own capping enzymes. In this communication, we characterize a novel single subunit mRNA capping enzyme from another NCLDV, Faustovirus. Unlike VCE, a heterodimeric enzyme that requires a small subunit to generate Cap-0 efficiently (Mao and Shuman 1994; Kyrieleis et al. 2014), the single-chain Faustovirus mRNA capping enzyme (FCE) generates Cap-0 from 5’ triphosphate RNA efficiently. FCE and the VCE large subunit (D1R) share a common catalytic domains organization. The two capping enzymes exhibit the same enzyme activities under similar optimal reaction conditions. Interestingly, FCE exhibits a higher specific activity in Cap-0 synthesis, operates under a broader active temperature range, and is more active on structured RNA.

## Results

### Identification and predicted features

Faustovirus mRNA capping enzyme (FCE; GenBank Accession AMN83561) is a single subunit RNA enzyme from Faustovirus, a giant virus that infects vermamoeba. AlphaFold2 prediction suggests that FCE is organized into three domains which are highly similar to the structure of corresponding domains of VCE D1R and the *Paramecium bursaria* chlorella virus (PBCV-1) GTase determined experimentally (Kyrieleis et al. 2014; Hillen et al. 2019; Håkansson et al. 1997) (Figure 1B). It shares very low sequence identity with the large subunit of VCE (D1R) (14.8%) and has no BLAST hits within poxviruses (data not shown), a double-stranded DNA virus family that vaccinia virus belongs to. However, manual inspection of the amino acids sequence showed that FCE possesses the conserved catalytic amino acids residues in capping enzymes from organisms such as Vaccinia virus, *S. cerevisiae*, *Paramecium bursaria* chlorella virus (PBCV-1) and human (Figure 1B). AlphaFold2 prediction suggests that FCE is organized into three domains which are highly similar to the structure of corresponding domains of VCE D1R and the PBCV-1 GTase determined experimentally (Kyrieleis et al. 2014; Hillen et al. 2019; Håkansson et al. 1997) (Figure 1B).

### Enzymatic activities

To verify that FCE generates the Cap-0 structure, we used purified FCE under standard VCE capping reaction conditions to cap a 25-nt RNA oligonucleotide containing a triphosphate group at the 5’ end. The reactions were then analyzed by nucleoside analysis using LC-MS. The results showed that FCE generates the same Cap-0 structure on RNA oligonucleotides with an adenosine or a guanosine as the initiating nucleotide (Suppl. Figure 2A).

We next verified the presence of the three individual enzyme activities. Using capillary electrophoresis as readout (Chan et al. 2022), we subjected FCE to conditions that facilitated the TPase activity, TPase and GTase activities, or all three activities. Using substrate RNA oligonucleotides containing a 5’ triphosphate group (ppp-), FCE generated a 5’ diphosphate end (pp-) in the absence of GTP and SAM, unmethylated G-capped 5’ end (Gppp-) in the presence of GTP only, and Cap-0 N7-methyl G-capped 5’ end (m7Gppp-) in the presence of both GTP and SAM (Suppl. Figure 2B). A time course of the FCE capping reaction in the presence of GTP and SAM showed that the conversion of 5’ ppp- to pp-, Gppp- and then m7Gppp- mirrored that of VCE reactions (Figure 2).

**Figure 2.**
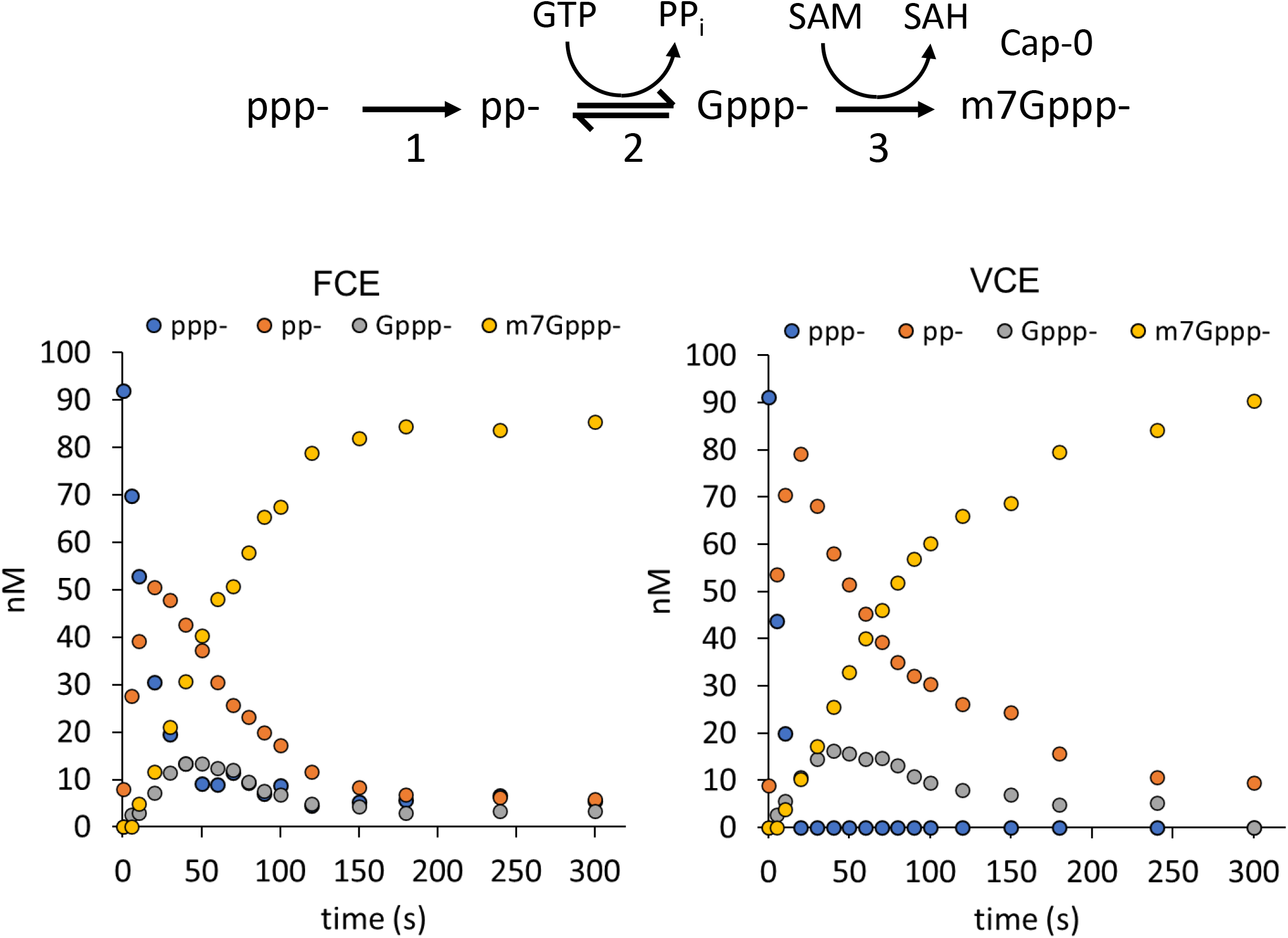
FCE and VCE generate the m7Gppp- (Cap-0) structure through the same pathway. Under identical conditions (1x RNA capping buffer, 0.5 mM GTP, 0.1 mM SAM, 5 nM enzyme at 25C), FCE and VCE underwent the same reaction pathway to generate the m7Gppp- cap from 5’ triphosphate. As the concentration of 5’ ppp-RNA (blue circles) decreased over time, the concentration of the TPase product pp- (orange circles) increased initially and decreased over time, followed by a similar trend with a smaller amplitude for the GTase product Gppp- (grey circles), and the increase in N7MTase product m7Gppp- (yellow circles).

To verify the active sites, we mutated FCE residues that correspond to important active site residues of VCE and compared them to wild-type (WT) FCE activity (Figure 1B, bottom panels). Assays were designed to interrogate each of the three enzyme activities quantitatively using substrate RNA containing a 5’ triphosphate, diphosphate, or unmethylated G-cap group. Results for WT FCE are shown in Figure 3A. The FCE TPase domain hydrolyzed over 90% of 5’ ppp-RNA to pp-RNA in Reaction 1 (in the absence of GTP and SAM). The GTase activity guanylylated ∼90% of 5’ pp-RNA to the Gppp- capped form in Reactions 2 (with GTP but no SAM). The FCE N7MTase activity methylated approximately 80% of Gppp-RNA to m7Gppp- in Reaction 3 (in the presence of SAM without GTP). The no enzyme controls showed that the ppp-RNA preparation contained a small amount of pp-RNA, the pp-RNA substrate preparation contained a small amount of ppp-RNA and the Gppp-RNA preparation contained a small amount of pp-RNA. These impurities were side products in the manufacturing of the oligonucleotide RNA molecules.

**Figure 3.**
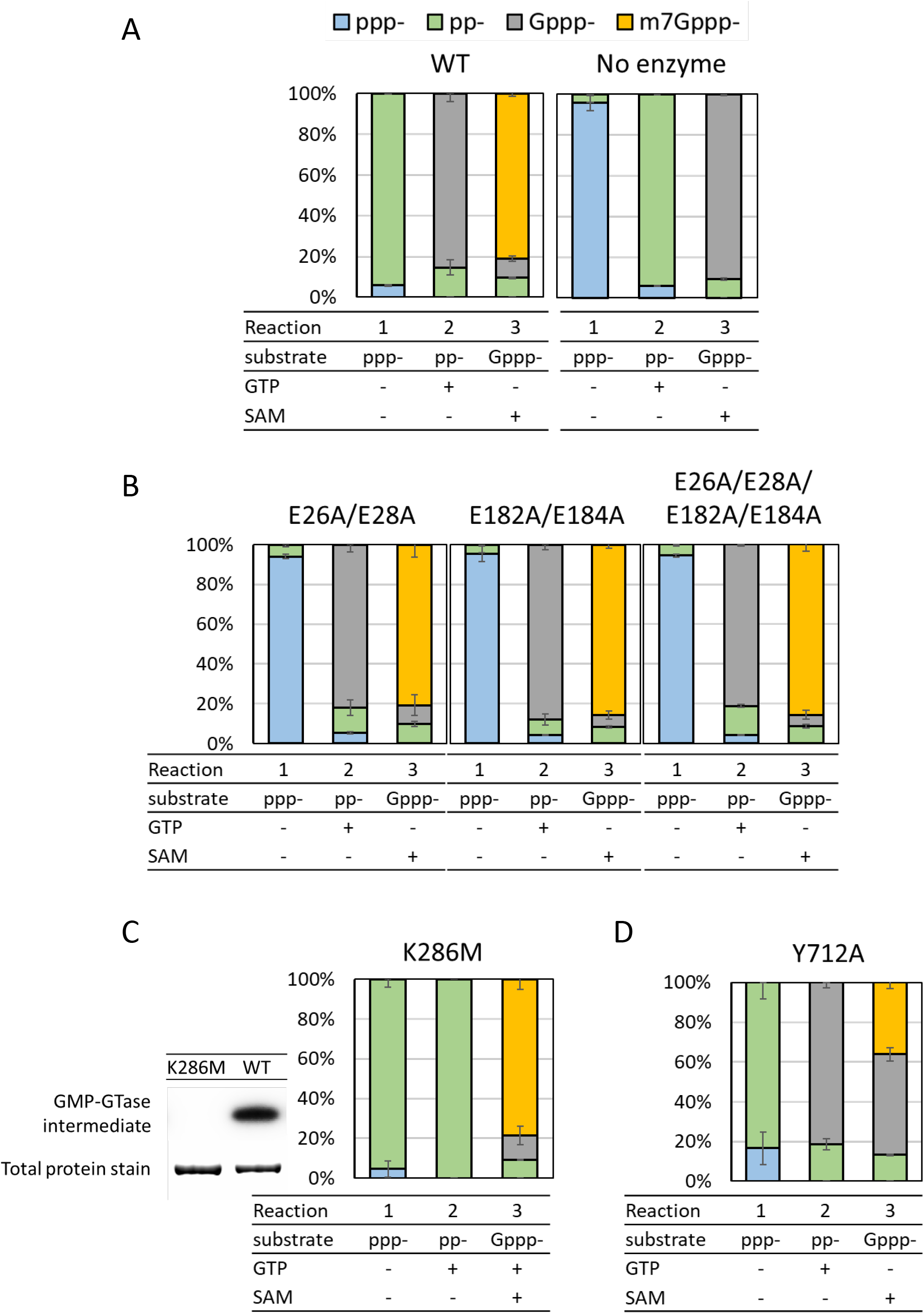
Catalytic amino acid residues of FCE. (A) Under standard conditions (see Materials and Methods), 2.5 nM of FCE WT enzyme converted ≥90% of ppp-RNA into pp-RNA in the absence of GTP and SAM (Reaction 1), ∼90% into Gppp- in the presence of GTP (Reaction 2) and ∼80% into m7Gppp- in the presence of GTP and SAM (Reaction 3). (B) Mutations of the Glu of the ELE motifs knocked out TPase activity but did not affect GTase or N7MTase activities. Mutants E26A/E28A, E182A/E184A and E26A/E28A/E182A/E184A exhibited the same trend. (C) Mutation of Lys286 of the conserved KTDG motif to Met eliminated the formation GMP-GTase covalent intermediate (left panel). The mutant K286M exhibited undetectable GTase activity (Reaction 2) but exhibited high level of TPase (Reaction 1) and N7MTase activity (Reaction 3). (D) Mutation of Tyr712 in the conserved SAM-binding motif to Ala significantly decreased the N7MTase activity (Reaction 3) but not TPase (Reaction 1) or GTase activity (Reaction 2).

### TPase

Alanine mutations were introduced to the two conserved metal-coordinating glutamic acids in the TPase domain ELE motifs (Figure 1B, left bottom panel). Double mutants E26A/E28A, E184A/E186A, and quadruple mutant E26A/E28A/E184A/E186A were generated and tested for the presence of the three enzyme activities. All three mutants exhibited a similar activity profile: the mutants were inactive in Reaction 1 but displayed similar activities in Reaction 2 and 3 (Figure 3B), showing that FCE contains the conserved ELE motifs required for its TPase reaction.

### GTase

FCE contains the conserved 282KTDG285 motif of adenosyl/guanylyl transferases where Lys282 is expected to form a covalent intermediate with GMP (Figure 1B, center bottom panel) (Soulière et al. 2008). We introduced a conservative K282M mutation and tested the enzyme’s ability to self guanylylate and guanylylate pp-RNA. Incubation with GTP in the presence of magnesium showed that mutant K282M is incapable of forming the covalent GMP intermediate compared to the WT enzyme (Figure 3C, left panel). Capping enzyme activity assays showed that the mutant was unable to convert pp-RNA into Gppp- in Reaction 2 but exhibited TPase and N7MTase activities in Reaction 1 and 3, respectively (Figure 3C, right panel), showing that Lys282 is the catalytic lysine residue required for the guanylyltransferase activity in FCE.

### N7MTase

The FCE N7MTase domain contains a conserved 707Q/NFAIHY712 SAM-binding motif, where the tyrosine residue in VCE D1R stacks with the adenosine ring of SAM and the rest of the motif spans the length of SAM binding pocket (Figure 1B, right bottom panel). Mutation of Tyr712 to Ala resulted in a significant decrease of N7MTase activity (Figure 3D, Reaction 3) but did not affect the TPase (Reaction 1) or GTase (Reaction 2) activities, showing the importance of Tyr712 in N7MTase activity in FCE.

### Biochemical characterization

FCE exhibits higher specific activity, defined as enzyme concentration to achieve 50% Cap-0 formation (Cap50), than VCE (Figure 4A). FCE capped 50% of the 5’ ppp-RNA substrate at ∼1 nM compared to VCE that capped 50% of the substrate at ∼2.5 nM at 37°C (Figure 4A). Similar to VCE, FCE does not exhibit preference towards RNA initiated with an adenosine or a guanosine (Figure 4B), the two most abundant 5’ nucleotides of eukaryotic and viral mRNA (Wang et al. 2019).

**Figure 4.**
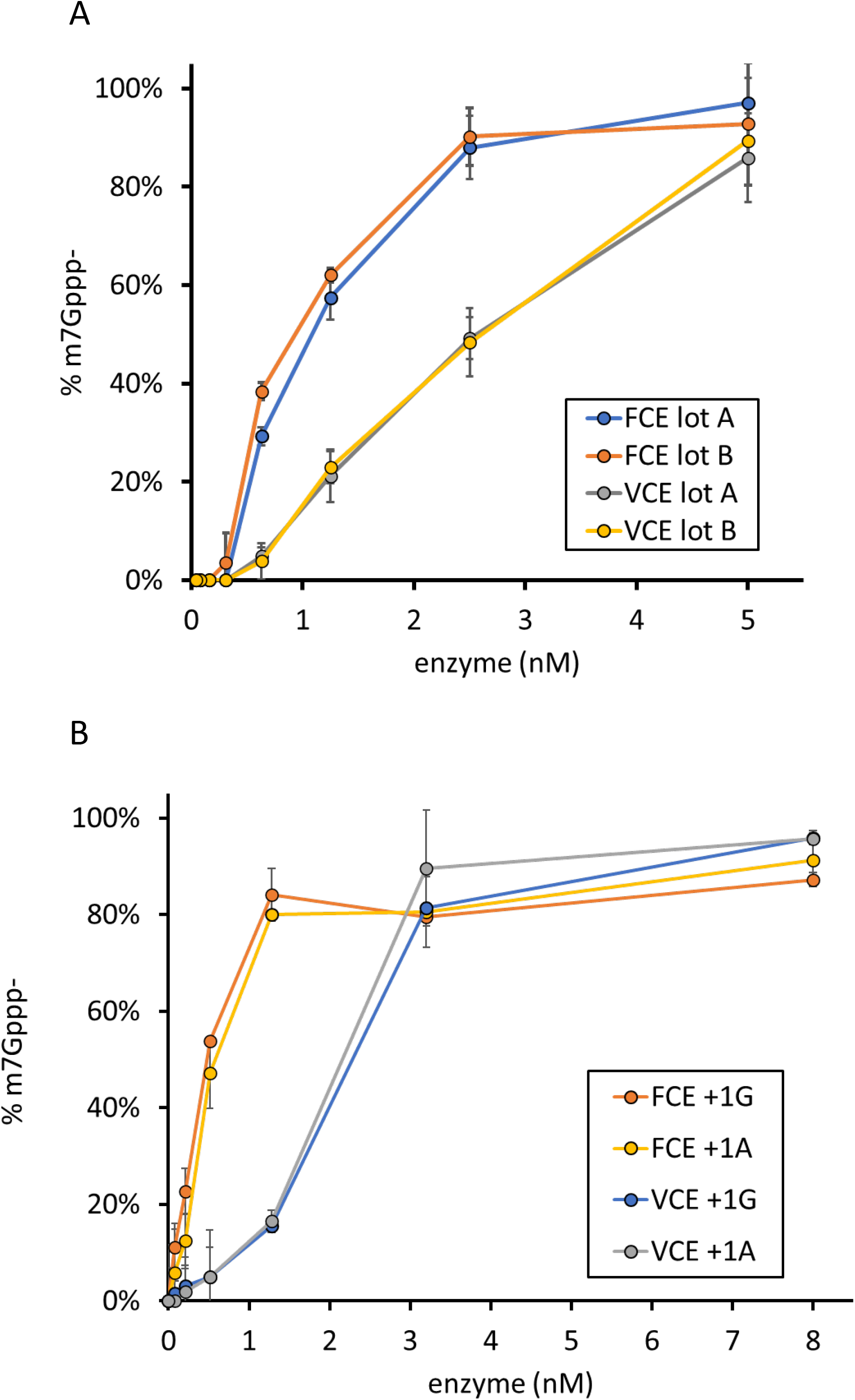
**(A)** FCE exhibits higher specific activity than VCE. RNA capping reactions were performed using decreasing concentrations of two preparations of FCE and VCE under standard conditions at 37°C for 30 min. FCE achieved 50% capping at ∼1 nM enzyme concentration; VCE achieved the same level of capping at ∼2.5 nM enzyme concentration. **(B)** FCE and VCE does not discriminate RNA initiating with guanosine or adenosine. RNA capping reactions were performed on 5’ triphosphate RNA oligonucleotide initiating with a guanosine (+1G) or with an adenosine (+1A) under standard conditions at 37C for 30 min. FCE and VCE exhibited similar level of capping activity on both +1G and +1A RNA. Data points are average value of triplicated experiments.

### pH

FCE exhibits highest Cap-0 generating activity at pH 7.0 – 8.0 (Figure 5A). Interestingly, our data showed that different enzyme activities were affected at either end of the pH range. We found that low pH (pH 6.0) impedes the TPase activity as the level of ppp-RNA remains high. High pH (pH 9.0) impedes the GTase and/or N7MTase activity as the level of pp-RNA remains high (Figure 5B). VCE exhibits a similar differential response.

**Figure 5.**
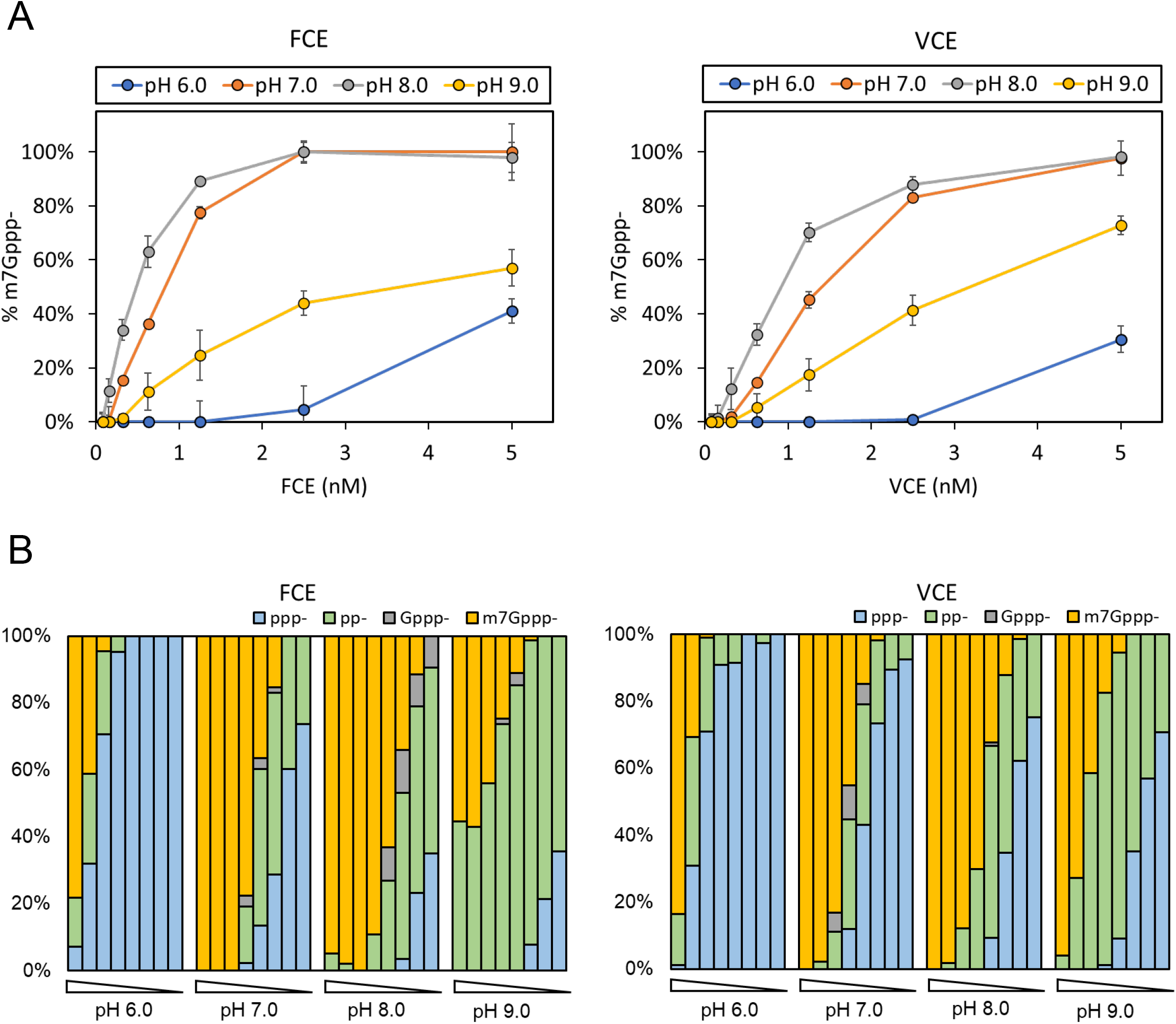
Effect of pH on capping activities. **(A)** FCE and VCE exhibit highest capping activity at pH 7.0 and pH 8.0. **(B)** Effect of pH on component enzyme activities. Both enzymes exhibit similar response outside of pH optimum. At pH 6.0, the accumulation of ppp- substrate indicates that the TPase activity of both enzymes are negatively affected. At pH 9.0, the accumulation of pp- product indicates that the forward GTase activity of both enzymes is negatively affected. Data points are average value of triplicated experiments.

### NaCl

Similar to VCE, FCE exhibits full Cap-0 capping activity up to 50 mM NaCl at low enzyme concentrations (Figure 6A). With 100 mM NaCl, both enzymes exhibited lower m7Gppp- generating activity but achieved close to 100% m7Gppp-RNA capping at 5 nM enzyme concentration. Interestingly, the TPase activity of FCE is more sensitive to increasing NaCl concentration than that of VCE, as indicated by the higher fraction of ppp-RNA at 200 mM NaCl and above in the FCE reactions (Figure 6B).

**Figure 6.**
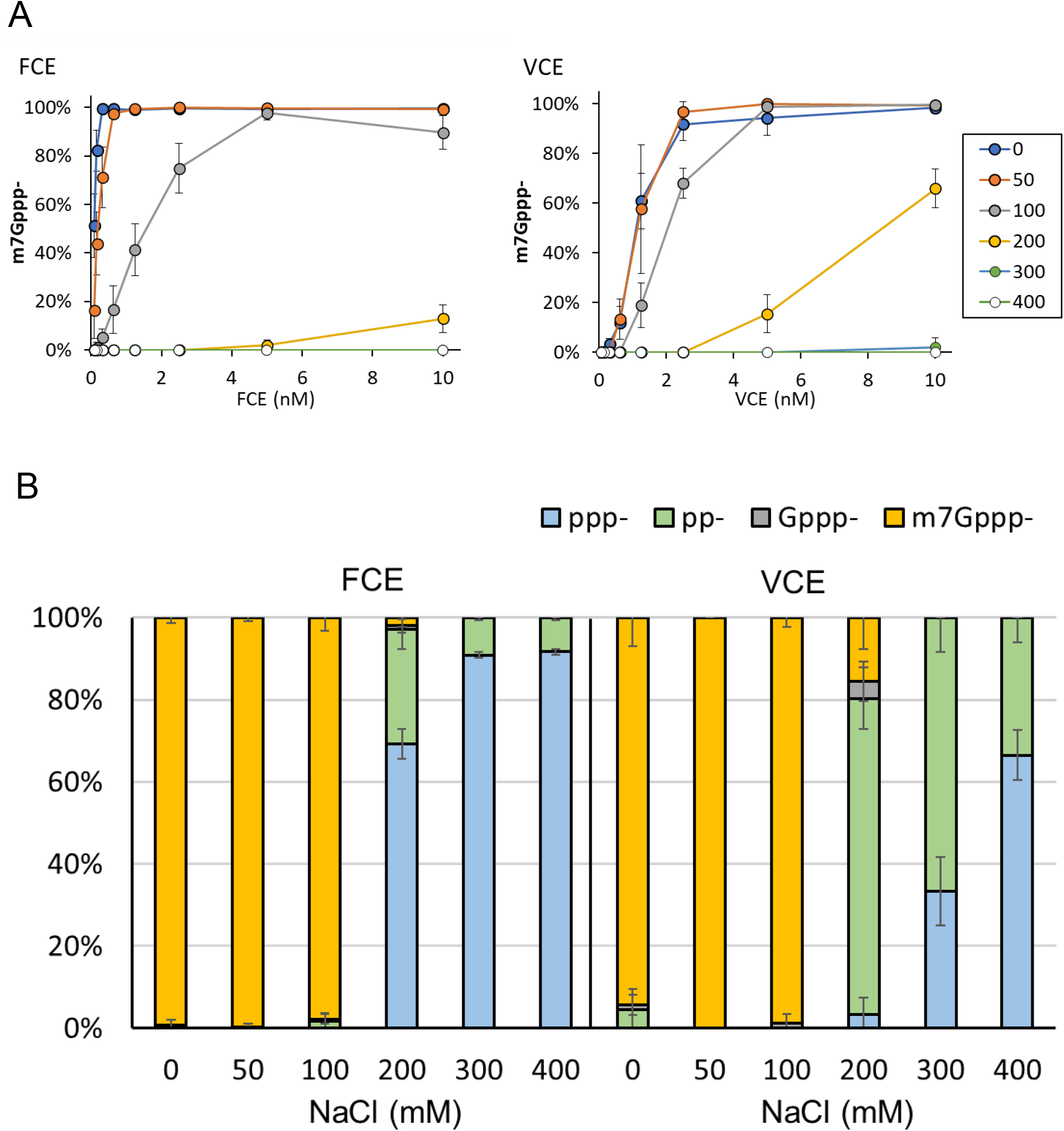
Effect of NaCl concentration on capping activities. **(A)** FCE and VCE exhibit highest capping activity in zero or 50 mM NaCl. At enzyme concentration of 5 nM or higher, both enzymes exhibit high capping activity in 100 mM NaCl. **(B)** Differential effect of NaCl on FCE and VCE. In 200 mM NaCl, the TPase activity of FCE decreased significantly whereas the GTase but not TPase activity of VCE was affected. The TPase activity of VCE started to be negatively impacted by 300 mM of NaCl. Data points are average value of triplicated experiments.

### MgCl2 concentration

Magnesium ion is required for the TPase and GTase activities. FCE and VCE exhibit the highest Cap-0 generating activity at 0.5-2 mM MgCl2 (Figure 7). The GTase activity of FCE was significantly impeded at 5 mM MgCl2, as evident by the accumulation of pp-RNA as reaction product. At the lower end of MgCl2 concentration, the TPase activity of FCE was greatly hampered at 0.1 mM MgCl2, as shown by the accumulation of ppp-RNA substrate, whereas the TPase activity of VCE was less affected under the same conditions. Our results show that FCE and VCE share similar optimal MgCl2 concentrations, but their component enzyme activities respond differently outside of this optimal range.

**Figure 7.**
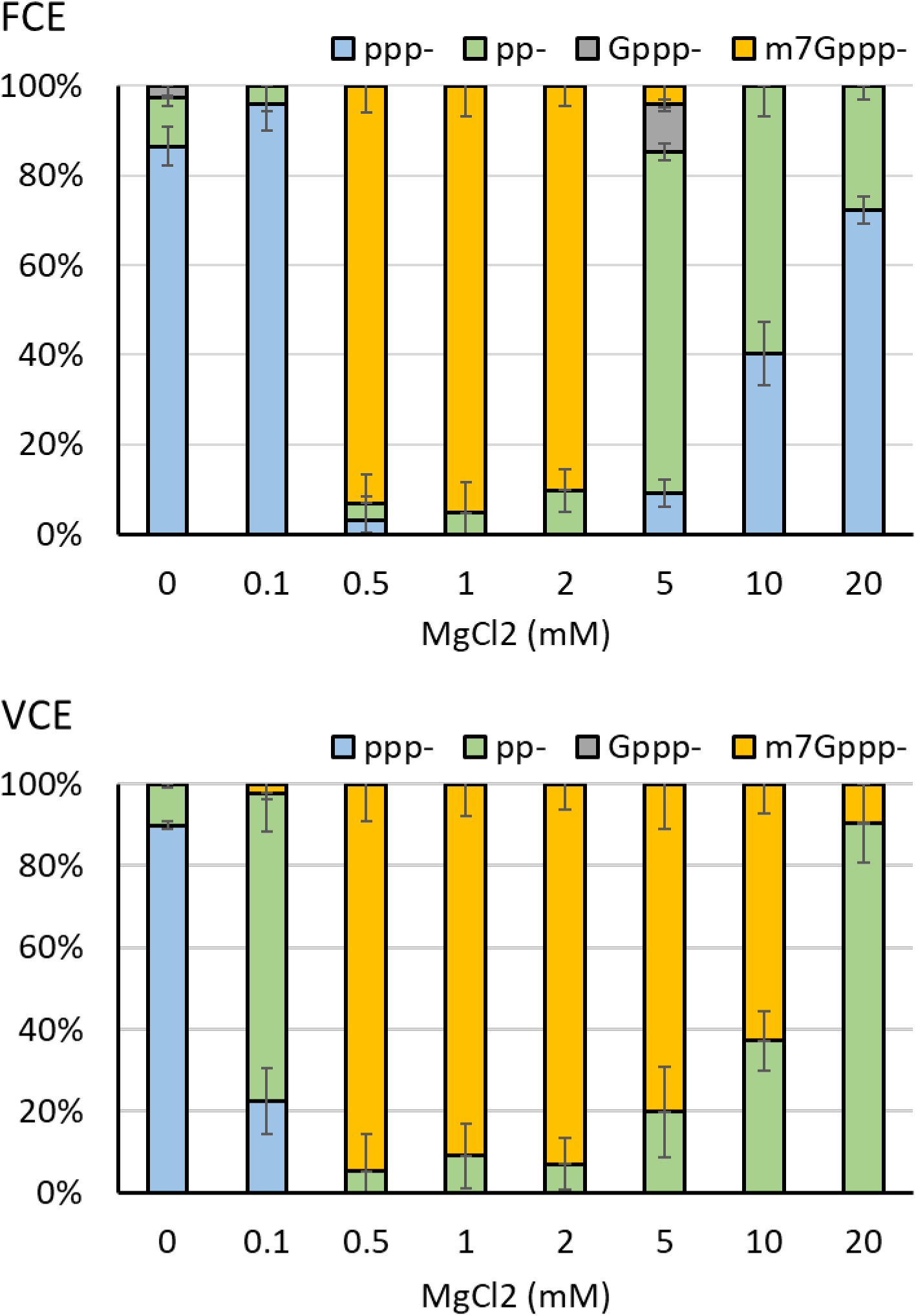
Differential effect of MgCl2 concentration on capping activities. FCE and VCE exhibit highest capping activity in 0.5 to 2 mM MgCl2 but enzyme activities of the two enzymes are differentially affected outside of the optimal MgCl2 concentration range. At 0.1 mM MgCl2, the TPase activity of FCE is impeded whereas in VCE the GTase activity was affected. For FCE, the GTase activity was negatively impacted at 5 mM MgCl2 with TPase started to be affected at 10 mM MgCl2. For VCE, the GTase activity started to be affected at 5 mM MgCl2 as pp-RNA accumulated. However, the TPase activity of VCE was not affected up to 20 mM MgCl2 due to the lack of ppp– accumulation. Data points are average value of triplicated experiments.

### Reaction temperature

FCE remains highly active Cap-0 incorporation between 25-55°C, compared to VCE that is most active between 35-45°C (Figure 8, blue and orange, respectively). Inspection of intermediate products shows that 5’ pp-RNA began to accumulate at 45°C and became the dominant product at 50°C and above for VCE, indicated a defect in TPase activity (Figure 8). For FCE, accumulation of 5’ pp-RNA product only began at 60°C, where a large fraction of reaction products was Gppp-RNA, suggesting that the N7MTase activity of FCE was compromised. On the other hand, the absence of Gppp-RNA product in the VCE high temperature reactions suggests that both the N7MTase and the GTase activities were compromised, or the N7MTase activity was compromised and the Gppp- products were rapidly converted back to pp- by the eversible GTase activity (Soulière et al. 2008). It should be noted that for both enzymes, very little to no ppp- substrate was detected at all the temperatures tested, showing that the TPase activity was active from 15°C to 60°C.

**Figure 8.**
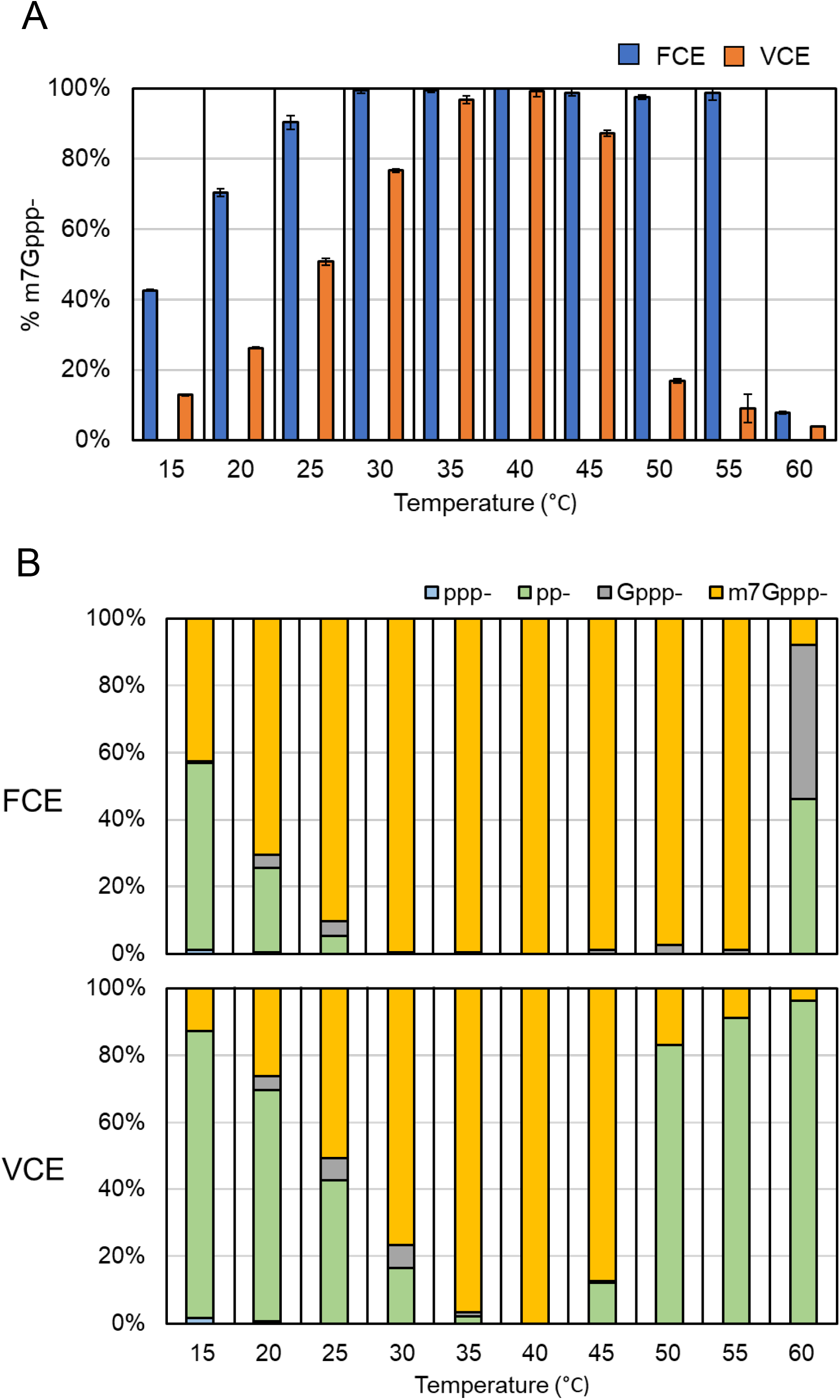
RNA capping activity of FCE and VCE across a temperature range. RNA capping assays were performed on 0.5 μM of a 20 nt poly(A) synthetic oligonucleotide with a 3’ FAM group using 5 nM of FCE or VCE under standard conditions (see Materials and Methods). **(A)** FCE generated more m7Gppp-cap at all temperatures tested. Data points are average of triplicated reactions. Error bars represent standard deviations. **(B)** A breakdown of the intermediate products across the temperature range.

### RNA Capping activity on structured 5’ ends

A series of RNA oligonucleotides (23-25 nt) were designed to investigate the effect of secondary structure and the availability of a free 5’ end on FCE and VCE capping activity (Figure 9). A relative low enzyme concentration (2 nM) was used to assess differences in capping activity of the two enzymes. Under these conditions, FCE generated more Cap-0 product than VCE on an RNA oligo with two unpaired nucleotides at the 5’ end within a range of hairpin structures with varying degree of thermodynamic stability (Oligos 1, 2 and 3). It generated less Cap-0 product on hairpin structure containing a one-nucleotide 5’ overhang or no overhang at a level similar to that of VCE (Oligos 4 and 5). No Cap-0 product was detected for either enzyme on model oligos with a stable stem-loop structure and recessive 5’ ends (Oligos 6 and 7). Therefore, FCE exhibit constraints of capping activity for stem-looped RNA structures and have a strong preference for free 5’ ends.

**Figure 9.**
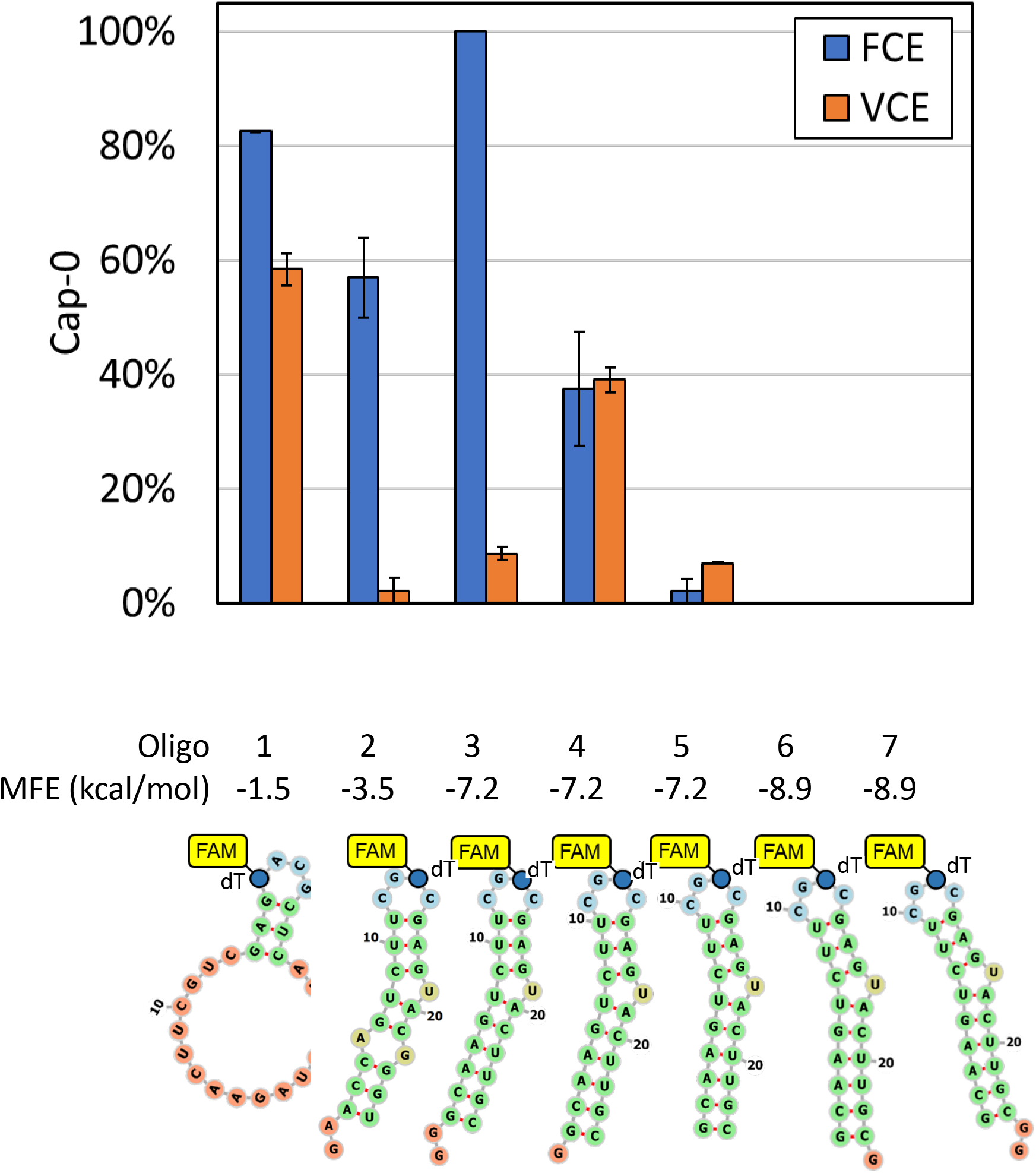
RNA capping activity of FCE and VCE on structured model RNA. Synthetic oligonucleotides designed to adopt hairpin structures with different folding energies and availability of 5’ triphosphate. All RNA oligos contains a FAM-dT for capillary electrophoresis readout. Capping reactions were carried out on 0.5 μM of the RNA oligos using 2 nM of FCE or VCE at 37°C under standard conditions (see Materials and Methods). Data points are average of triplicated reactions. Error bars represent standard deviations.

### RNA Capping activity on *in vitro* transcripts

To compare the RNA capping activity of FCE and VCE on *in vitro* transcripts, 20 nM FCE or VCE was used to cap 3.5 μM (2 μg/μL) of *FLuc* transcripts (1.8 kb) containing the 5’ sequence of the first 50 nt from human beta globin and synthetic transcripts derived from pRNA21, *FLuc*, and Comirnaty in 100 μL reactions for 1 hour at 37°C. The extent of cap incorporation was measured using targeted RNase H cleavage and LC-MS (Chan et al. 2022). Overall, transcripts containing different 5’ sequences were capped to different extent by both enzymes, and FCE generated a higher percentage of m7Gppp- cap than VCE in the *in vitro* transcripts tested (Figure 10).

**Figure 10.**
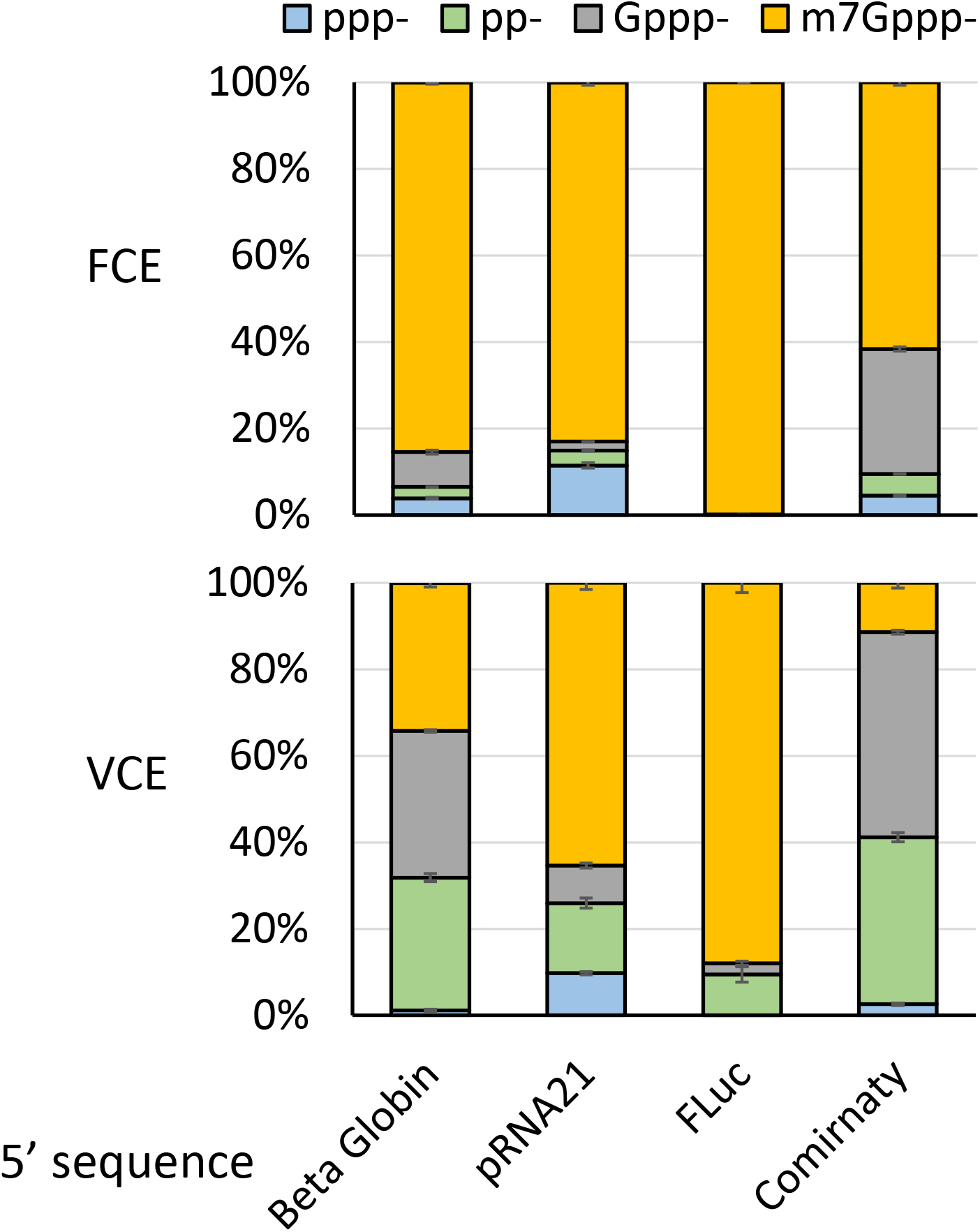
RNA capping activity of FCE and VCE on *FLuc* transcript (1.8 kb) containing 5’ sequence derived from the first 50 nt of human beta globin variant 1, synthetic 5’ UTR of pRNA21, synthetic 5’ UTR of p*FLuc*, and Comirnaty, respectively. 20 nM of FCE or VCE was used to cap 3.5 μM of the *in vitro* transcripts for 1 hour at 37°C. Following capping reactions, mRNA capping was measured using targeted RNase H cleavage and LC-MS. Datapoints are average of triplicated experiments.

### Phylogenetic analysis

Vaccinia virus and Faustovirus belong to a monophyletic group of eukaryotic double-stranded DNA viruses called the nucleocytoplasmic large DNA viruses (NCLDV). As a group of dsDNA viruses that are diverse in genome size (ranges from 100 kb to 2.5 Mb), virion morphology and host range that includes amoeba, unicellular algae, insects and humans, NCLDVs only share three conserved genes – genes that code for a family B DNA polymerase, a primase-helicase and a predicted transcription factor required for late transcription in poxvirus VLTF3 (Koonin and Yutin 2019). However, NCLDVs share a core set of genes necessary for cytoplasmic viral transcription and replication, and capsid formation (Koonin and Yutin 2019). Cytoplasmic viruses do not have access to the host transcription and RNA capping machineries in the nucleus and face selective pressure to maintain an effective transcription and RNA capping mechanism. Hence, the mRNA capping machinery is broadly conserved among the NCLDV, with the exception of those that have adapted nuclear transcription and replication (Allen et al. 2006; Chinchar et al. 2017).

To examine the conservation FCE and survey the diversity of RNA capping enzymes within the virosphere, viral genomes were searched for the presence of the cap N7MTase domain (Pfam PF03291). Eighty-three genomes, all belonging to the NCLDV, were identified and clustered based on proteome similarity using the program vContact2 (Figure 11A) (Bin Jang et al. 2019). The amino acid sequences of representative members of the identified capping enzyme genes were aligned, and their sequence similarity is represented in the form of a phylogenetic tree (Figure 11B). These capping enzyme genes were then mapped to the proteome similarity networks. We found that phylogenetic analysis of the capping enzymes is largely consistent with proteome similarity networks consistent with the notion that broad conservation and correlation with proteome-similarity highlights the importance of capping enzyme to the lifestyles of these viruses (Koonin and Yutin 2019)(Figure 11A). The capping enzyme from *Faustoviruses* are most similar to those from African swine fever viruses and Pacmanviruses (Figure 11B) while the three families of viruses cluster in the same proteome similarity network (blue dots, Figure 11A). Most of the clades, except Panodoraviruses (light orange) and Phycodnaviruses (dark orange) possess putative RNA capping enzymes as multifunctional proteins containing all three enzyme activities. Poxvirus capping enzymes are the only group of enzymes that possess an obvious activation subunit (D12L of Vaccinia virus) and form a distinct branch on the phylogenetic tree (green dots) (Figure 11B). Similar to a previous study, the poxviruses form a distinct group in the our proteome similarity networks analysis (Figure 11A) (Guglielmini et al. 2019). Interestingly, Pandoravirus and Phycodnavirus capping enzymes that have separated TPase, MTase and GTase genes are clustered under a common ancestor distinct from the multifunctional capping enzymes (Figure 11B), even though the two groups of NCLDVs are clustered to distinct proteome similarity networks.

**Figure 11.**
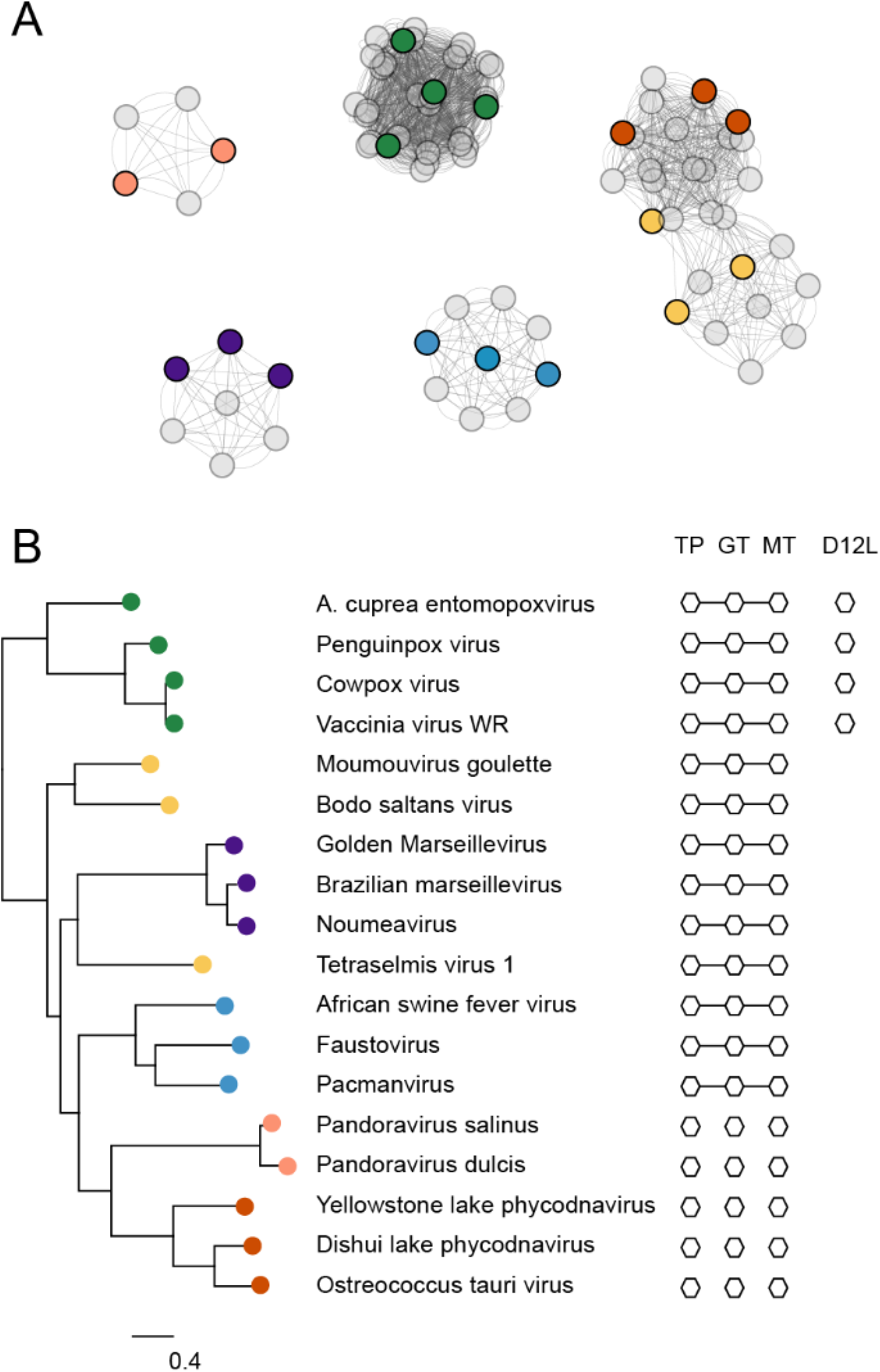
**(A)** The CheckV database of complete viral genomes was searched for viruses which contain a cap M7MTase domain. The 83 identified viruses were clustered by proteome similarity using the vContact2 program. **(B)** A subset of the 83 capping enzymes were selected to represent the diversity of the data set, aligned and used to construct a phylogenetic tree. The tree is built from a MAFFT alignment of the N7MTase domains; for viruses which contain separate genes from TPase, GTase and N7MTase domains, only the N7MTase protein was used. The viruses were mapped to the clustering in (A) (color dots), showing that the clustering based on whole proteome is consistent with the phylogeny of the cap N7MTase genes. On the right, the presence of TPase (TP), GTase (GT) and N7MTase (MT) domains, as well as regulatory subunit homologs (D12L) of Vaccinia virus, within the genomes is denoted. Domains attached by a solid line are found within one gene.

## Discussions

In this communication, we presented biochemical and enzymatic characterization of a novel single subunit RNA capping enzyme from Faustovirus. Faustovirus has been shown to infect *Vermamoeba vermiformis*, an amoeba that has been found in human-associated environments such as hospital water networks, drinking water, and human stool samples (Pagnier et al. 2015; Bradbury 2014; Delafont et al. 2018; Hajialilo et al. 2015; Rolland et al. 2019). As a giant virus (NCLDV that have a genome size of 400 kb or larger) with a 460 kb dsDNA genome, our phylogenetic analysis based on two universally conserved genes (Family B DNA polymerase and major capsid protein), as well as previously published analysis (Koonin and Yutin 2019), suggests that Faustovirus is more closely related to Asfavirus (genome size of 170-190 kb) than giant viruses of similar genome sizes (such as Cedratvirus of 590 kb or Marseillevirus of 360 kb) (Koonin and Yutin 2019). Amino acid sequence analysis also showed that FCE is more closely related to Asfavirus RNA capping enzyme (Figure 11B). Our phylogenetic analysis suggests that NCLDVs derive their RNA capping enzymes from a common ancestral enzyme which is a single-subunit protein with multiple-activities. It is also likely that the functional dependence of an activation subunit of poxvirus RNA capping enzymes (e.g., Vaccinia virus D12L) occurred after the speciation event that produced the ancestors of the Poxviridae and Asfarviridae such that poxvirus RNA capping enzymes remains the only NCLDV capping enzymes that requires an activation subunit.

Similar to NCLDV RNA capping enzymes, RNA capping enzymes in eukaryotes adopt a diverse configuration. Fungal capping enzymes are composed of separate TPase, GTase and N7MTase proteins (e.g., Cet1, Ceg1 and Abd1 of *S. cerevisiae*), whereas higher eukaryotes have a bifunction TPase and GTase protein (e.g., RNGTT of humans and Drosophila that contains a Tyrosine protein phosphatase domain for TPase activity) and a N7MTase (e.g., RNMT of humans). It is not clear whether the separation of these enzyme activities on individual proteins contributes to any biological functions. But it is fascinating that the modulate nature of RNA capping enzymes allows the evolution of so many forms throughout eukaryotes and their viruses (see (Ramanathan et al. 2016) for an overview of the RNA capping machineries of RNA viruses) while each being functionally adapted to the organisms as an essential component of the transcription-translation machinery.

It is fortuitous that FCE exhibits higher specific activity and broader temperature range than VCE. Such properties can be desirable in mRNA manufacturing applications and may enable major improvements in mRNA manufacturing workflows. As with most enzymes, *in vitro* activities of FCE can vary based on reaction conditions such as enzyme, substrate and reagent concentrations, supplementation of additives, temperature, and accessibility of the substrate. For RNA capping enzymes, the true substrate is the 5’ triphosphate group of newly synthesized RNA. Our data and previously published results showed that the secondary structure of the 5’ region plays a significant role in the “cappability” of the RNA by capping enzymes (Fig 9) (Fuchs et al. 2016). We further demonstrated that the 5’ sequence of *in vitro* transcripts can have a significant impact to the “cappability” of the transcripts (Figure 10). Therefore, exploring the sequence space for highly “cappable” 5’ ends, in addition to *in vivo* expression levels (Sample et al. 2019; Jia et al. 2020; Leppek et al. 2022), would be beneficial for manufacturing of high-quality mRNA that offers high translation efficiency and low immunogenicity for therapeutic applications.

## Materials and Methods

### Enzymes and RNA substrates

Faustovirus RNA capping enzyme (FCE) and Vaccinia RNA capping enzyme (VCE) are obtained from New England Biolabs, Inc. FCE mutant constructs were generated by GenScript (Piscataway, NJ) by site-directed mutagenesis. His-tagged FCE mutant proteins were expressed from *E. coli* BL21(DE3)star and purified by Ni-column chromatography and size-exclusion chromatography by GenScript to over 90% homogeneity (determined by Coomassie blue-stained SDS-PAGE). RNA oligonucleotides containing 5’ triphosphate, 5’ diphosphate, 5’ Gppp- cap and 5’ m7Gppp- cap are synthesized by Bio-synthesis, Inc. (Lewisville, TX).

### Enzyme assays

Unless specified otherwise, RNA capping enzyme assays were performed in 10 μL of 1x RNA capping buffer (New England Biolabs, Inc.) containing 0.5 μM substrate RNA, 0.1 mM SAM, 0.5 mM GTP, 1 mM DTT, and specified concentration of capping enzymes. The reactions were carried out in a thermocycler for 30 minutes at a specific temperature and quenched by adding equal volume of a 2x stop solution containing 20 mM EDTA and 2% SDS. The quenched reactions were diluted for capillary electrophoresis on an Applied Biosystems 3730xl Genetic Analyzer (96 capillary array) using POP-7 polymer and GenScan 120 LIZ dye Size Standard (Chan et al. 2022; Wulf et al. 2019; Greenough et al. 2016). Peak detection and quantification were performed using the Peak Scanner software v.1.0 (Thermo Fisher Scientific) and an in-house data analysis suite. Validation of the capillary electrophoresis platform for RNA cap analysis has been published (Chan et al. 2022).

### Verification of the identity of 5’ cap

RNA oligonucleotides starting with a +1G (pppG-25mer) and +1A (pppA-25mer) were capped using FCE under standard conditions: 0.5 μM substrate RNA, 0.1 mM SAM, 0.5 mM GTP, 1 mM DTT, 100 nM FCE in 10 μL of 1x RNA capping buffer for 30 min at 37°C. The reactions were treated by Nucleoside Digestion Mix (New England Biolabs, Inc.) to release the nucleosides and the dinucleotide cap structure (m7GpppG or m7GpppA). Specifically, 4 μL of 10x Nucleoside Digestion Mix reaction buffer, 24 μL of nuclease-free water and 2 μL of the Nuclease Digestion Mix were added to 10 μL of the capping reactions. After the reactions were incubated at 37°C for 1 h, the samples were subjected to LC-MS nucleoside analysis using an Agilent 1200 Series LC-MS System equipped with a G1315D diode array detector and a 6120 Single Quadrupole Mass Detector in both positive (+ESI) and negative (-ESI) electrospray ionization modes. LC was performed on a Waters Atlantis T3 column (4.6 × 150 mm, 3 μm) with a gradient mobile phase consisting of aqueous ammonium acetate (10 mM, pH 4.5) and methanol. The identity of each peak was confirmed by MS. The relative abundance of nucleosides was determined by comparing the respective peak area at 260 nm or their respective UV absorption maxima to that of cytidine in each sample.

### Self-guanylylation assay

In a 20 μL reaction containing 1x capping buffer, 100 nM of WT FCE or mutant K286M was incubated with 1 μM of ^32^P-α-GTP (Perkin Elmer) in a 20 μL reaction containing 1x RNA capping buffer at room temperature for 1 minute. The reactions were quenched by adding 10 μL of 3X SDS loading dye (New England Biolabs, Inc.). The quenched reactions were analyzed by SDS-PAGE on a 10-20% Tris-Glycine SDS polyacrylamide mini-gel (ThermoFisher). The presence of the GMP-GTase covalent intermediate was visualized by autoradiography using a Phosphor Storage Screen (Cytiva) and the Amersham Typhoon RGB scanner (Cytiva).

### Capping activity assay on 5’ secondary structures

A series of RNA oligonucleotides was designed to possess stem-loop structures of different stability and different accessibility to the 5’ triphosphate group (Figure 9). Oligo 1 is expected to form a short 3-bp stem with an unstructured 5’ end. Oligo 2 is expected to form a longer stem structure with mismatches such that the theoretical minimum free energy of unfolding (MFE) is a moderate -3.5 kcal/mol with the first two nucleotides free from base-pairing. Oligo 3 is also expected to have two free nucleotides at the 5’ end but adopt a more stable stem-loop structure with a MFE of -7.2 kcal/mol. Oligo 4 and 5 and have the same secondary structure but are expected to have one free nucleotide at the 5’ end and a fully base-paired 5’ end, respectively. Oligos 6 and 7 are expected to have a slightly more stable stem-loop structure with MFE of -8.9 kcal/mol and one-base and two-base recessive 5’ ends, respectively. The RNA oligonucleotides contained a FAM-conjugated dU in the loop region to facilitate readout using capillary electrophoresis. To facilitate differentiating capping activity of FCE and VCE on the structured RNA oligonucleotides, a lower concentration of enzymes (2 nM) was used in the assay with 0.5 μM of the structured RNA oligos at 37°C for 30 minutes under standard conditions. The reactions were quenched and analyzed by capillary electrophoresis as described above.

### Phylogenetic analysis

The CheckV database of viral genomes (Nayfach et al. 2021) was searched for viruses encoding capping enzymes using the hmmer program (Wheeler and Eddy 2013) and the PFam profile for the poxvirus N7MTase domain (Pfam: PF03291). The resulting viral genomes were all members of the NCLDV and these were clustered on the basis of proteome similarity using vContact2 (Bin Jang et al. 2019). Associated capping enzymes were aligned with MAFFT (Madeira et al. 2022) using the G-INS-I algorithm and used as an input for FastTree (Price et al. 2010) with default parameters to produce a phylogenetic tree which was rooted according to (Guglielmini et al. 2019). The initial alignment of capping enzymes was used together with the VCE crystal structure (Kyrieleis et al. 2014) and predicted structures (Mirdita et al. 2022) to define boundaries between TPase, GTase and MTase domains. Profiles for these domains were constructed and used to identify distant homologs of the poxvirus capping enzyme domains in viral genomes for which they occur in separate genes. To identify Vaccinia D12L homologs, NCLDV translated proteomes were searched with the existing profile hidden Markov model (PFAM: PF03341).

## Acknowledgements

We would like to thank the NEB sequencing core for the of capillary electrophoresis analysis as well as all the facilities personnel for daily support of operation. This work is supported by New England Biolabs, Inc.

