## Supplemental Materials for "Biochemical characterization of mRNA capping enzyme from Faustovirus"

**Table 1.** Reference sequences used in this study.

| Protein | GenBank accession number |
|---|---|
| FCE | MN83561.1 |
| VCE D1R | AAO89385.1 |
| VCE D12L | AAO89396.1 |
| ASFV RNA capping enzyme | QZK26801.1 |
| hRNGTT | NP_003791.3 |
| hRNMT | NP_001365064.1 |
| Cet1 | KZV07283.1 |
| Ceg1 | CAA60705.1 |
| Abd1 | AAA34383.1 |
| *Paramecium bursaria* Chlorella virus GTase | AAC96471.1 |

**Table 2.** RNA oligonucleotides used in this study.

| Name | Sequence |
|---|---|
| pppG-25mer | pppGUAGAACUUCGUCGAGUACGCUCAA |
| pppA-25mer | pppAUAGAACUUCGUCGAGUACGCUCAA |
| Oligo-01 | pppGUAGAACUUCGUCGAG[FAM-dT]ACGCUCAA |
| Oligo-02 | pppGAACCAGUCUUCG[FAM-dT]CGAGUACGGGU |
| Oligo-03 | pppGGGCAAGUCUUCG[FAM-dT]CGAGUACUUGC |
| Oligo-04 | pppGGCAAGUCUUCG[FAM-dT]CGAGUACUUGC |
| Oligo-05 | pppGCAAGUCUUCG[FAM-dT]CGAGUACUUGC |
| Oligo-06 | pppGCAAGUCUUCG[FAM-dT]CGAGUACUUGCGG |
| Oligo-07 | pppGCAAGUCUUCG[FAM-dT]CGAGUACUUGCG |

**Supplemental Figures Legends**

**Suppl. Fig. 1 (A)** Structures of multi-domain capping enzymes from Faustovirus and Vaccinia. The FCE structure was predicted using ColabFold while the VCE D1R and D12L subunits were extracted from the CryoEM co-transcriptional capping complex (PDB ID 6RIE). Domain organizations were annotated. **(B)** FCE secondary structure prediction, active site residues and conserved active site motifs.

**Suppl. Fig. 2 (A)** Verification of m7Gppp- caps generated by FCE. Synthetic RNA containing 25 nucleotides with a guanosine or an adenosine as the first nucleotide were treated with or without FCE followed by treatment with the nucleoside digestion mix containing Nuclease P1. CIP and DNase I. The digested reactions were subjected to UPLC analysis. The area under the designated peaks were used to calculate the fraction of indicated species after correcting for their extinction coefficients. The values were normalized to the that for cytidines. **(B)** Reaction products of FCE component enzyme activities. Reactions were carried out to allow for the designated activities to take place. The reactions were quenched and analyzed by capillary electrophoresis. Rxn 1 is a no enzyme control where the major peak in the electropherogram is the unreacted substrate 5' triphosphate RNA. In the absence of GTP and SAM, FCE converted the 5' triphosphate into diphosphate in Rxn 2. When supplemented with GTP, FCE converted the 5' triphosphate of the substrate RNA into the Gppp- cap structure (Rxn 3). In the presence of GTP and SAM, the 5' triphosphate group was converted to the m7Gppp- cap structure.

### Supplemental figures

Suppl. Fig. 1

A

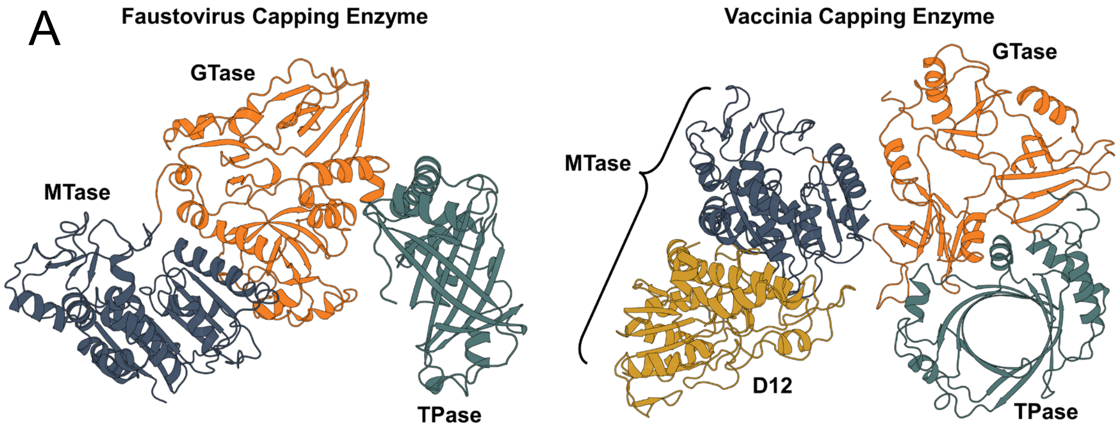


B

```
          HHHHHHHHHHH    SSSSSS    HHHHHHHHHHHH   SSSSSSSSSS  SSSSSSSSS  SSSSSSS
MAKRLQRCQDVNQVCEIYNSKGGIGELELRFDKLPQNLFAGVFDKLKPDGEIQTTMRVSNRDGVAREITFGGGVKTNEIF
         |         |         |         |         |         |         |         |
        10        20        30        40        50        60        70        80

SSSSSSSSSSS     SSSSSSSSSS           SSSSSSSSSS      SSSSSSSSSSSSSHHHH  HHHHHHHH
VKKQNICVFDVVDIFSYKVAVSTEETVVEKPTMETTAGVRFKIRLSVEDVVKDWRIDLTAVKTAELGKIAQHTASIVQRT
         |         |         |         |         |         |         |         |
        90       100       110       120       130       140       150       160

    HHH  HHHHHHH  SSSSSSSSS       HHHHHHHHHHHHHHHHHHHHHH HHHHHH HHHHHHHHHHHHH  H
FPDNLLKLTGAEVAKLAADSYELELEYTGKSPATNEKVNVAAKYAVELLSSVRNANSTAAASFGESVSDLCRVAKIIHTH
         |         |         |         |         |         |         |         |
       170       180       190       200       210       220       230       240

HHHHHHHH   HHHH        HHHHHH      SSSSSSS   SSSSSSSS  SSSSS    SSSSS      SSSSS
EYANVVCRTPSFKMLLPQVVSLTKSSYYGGLYPPENLWLAGKTDGVRALVVCEDGVAKVITAESVDITHGVCSATTILDC
         |         |         |         |         |         |         |         |
       250       260       270       280       290       300       310       320

SSS   SSSSSSSSSSS    HHH  HHHHHH           SSSS   SSS    HHHHHHH      SSSSSS
ELNVDAKILYVFDVIISNNTQVYTQPFSTRITTDISDIKIDGYKIEMKPFVKVVKADEATFKSAYKAPHNEGLIMIEDGA
         |         |         |         |         |         |         |         |
       330       340       350       360       370       380       390       400

 HHH  SSSS       SSSSSSS  HHH           SSSSSSSSSSHHHHHHH        HHHH         SS
AYAATKTYKWKPLSHNTIDFLIKACPKQLINVDPYKPRAGYKLWLLFTTISLDQQRELGIEFIPAWKILFTDINMFGSRV
         |         |         |         |         |         |         |         |
       410       420       430       440       450       460       470       480

SSS          SSSSS       SSSSSSSSSS      SSSSSSS HHHH         HHHHHHHHHHHH    HH
PIQFQPAINPLAYVCYLPEDVNVNDGDIVEMRAVDGYDTIPKWELVRSRNDRKNEPGFYGNNYKIASDIYLNYIDVFHFE
         |         |         |         |         |         |         |         |
       490       500       510       520       530       540       550       560

HHH            HHHHHHHHHHHHHHHHHHHHH     SSSSS       HHHHHHH   SSSSSS  HHHHHHHHH
DLYKYNPGYFEKNKSDIYVAPNKYRRYLIKSLFGRYLRDAKWVIDAAAGRGADLHLYKAECVEHLLAIDIDPTAISELVR
         |         |         |         |         |         |         |         |
       570       580       590       600       610       620       630       640

HHHHHHH                        SSSSSS      HHHHHHHHHHH        SSSSS  HHHH   HHHH
RRNEITGYNKSHRGGRNMHSHRGQSHCAKSTSLHALVADLRENPDVLIPKIIQSRPHERCYDAIVINFAIHYLCDTDEHI
         |         |         |         |         |         |         |         |
       650       660       670       680       690       700       710       720

HHHHHHHHHH   SSSSSSSSS HHHHHHHHHHH    SSSSSSS            SSSSSS            SSSS
RDFLITVSRLLAPNGVFIFTTMDGESIVKLLADHKVRPGEAWTIHTGDVNSPDSTVPKYSIRRLYDSDKLTKTGQQIEVL
         |         |         |         |         |         |         |         |
       730       740       750       760       770       780       790       800

       SSSS   HHHHHHHHHHH SSSSSSSSHHHHHHHHHHH HHHHHH  HHHHHHHHSSSSSSSS
LPMSGEMKAEPLCNIKNIISMARKMGLDLVESANFSVLYEAYARDYPDIYARMTPDDKLYNDLHTYAVFKRKK
         |         |         |         |         |         |         |
       810       820       830       840       850       860       870
```

Suppl. Fig. 2A

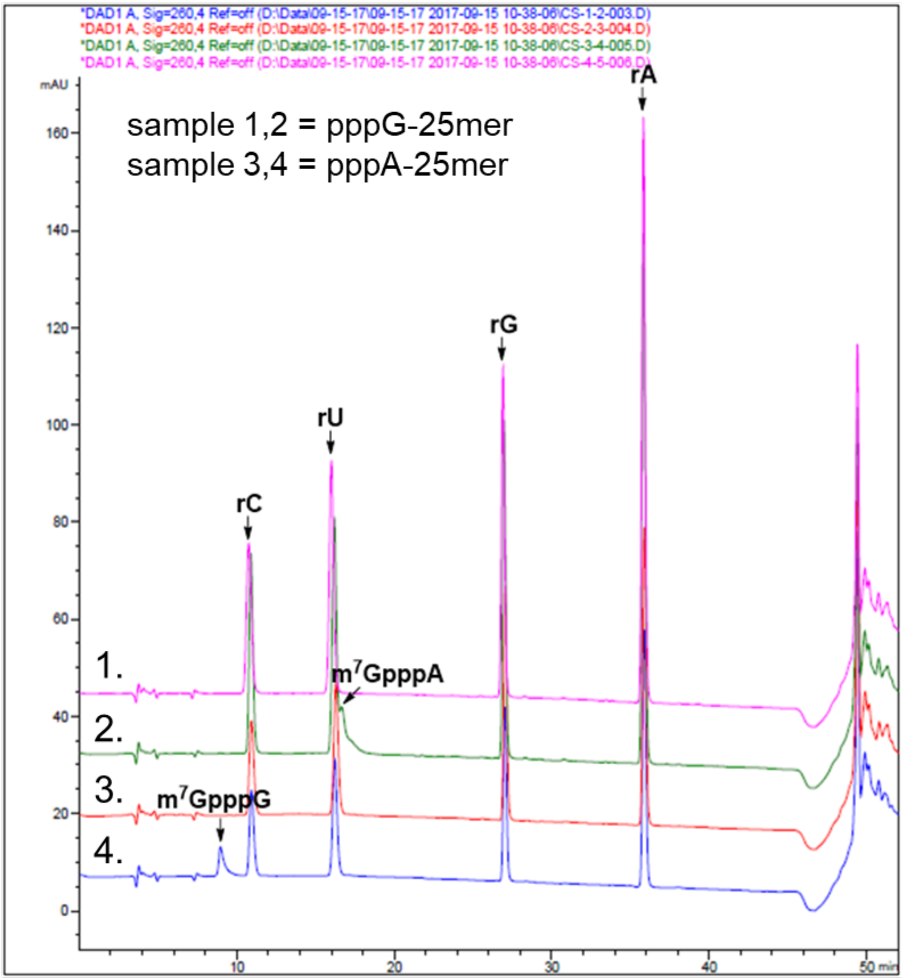


| | FCE | m7GpppG | C | U | m7GpppA | G | A |
|---|---|---|---|---|---|---|---|
| Sample 1 | + | 1.055 | 6.000 | 6.023 | 0.000 | 5.331 | 6.637 |
| Expected | | 1 | 6 | 6 | 0 | 5 | 7 |
| Sample 2 | - | 0.000 | 6.000 | 5.981 | 0.000 | 6.169 | 6.571 |
| Expected | | 0 | 6 | 6 | 0 | 6 | 7 |
| Sample 3 | + | 0.000 | 6.000 | 6.081 | 0.784 | 4.973 | 6.652 |
| Expected | | 0 | 6 | 6 | 1 | 5 | 7 |
| Sample 4 | - | 0.000 | 6.000 | 5.985 | 0.000 | 5.011 | 7.481 |
| Expected | | 0 | 6 | 6 | 0 | 5 | 8 |

Suppl. Fig. 2B

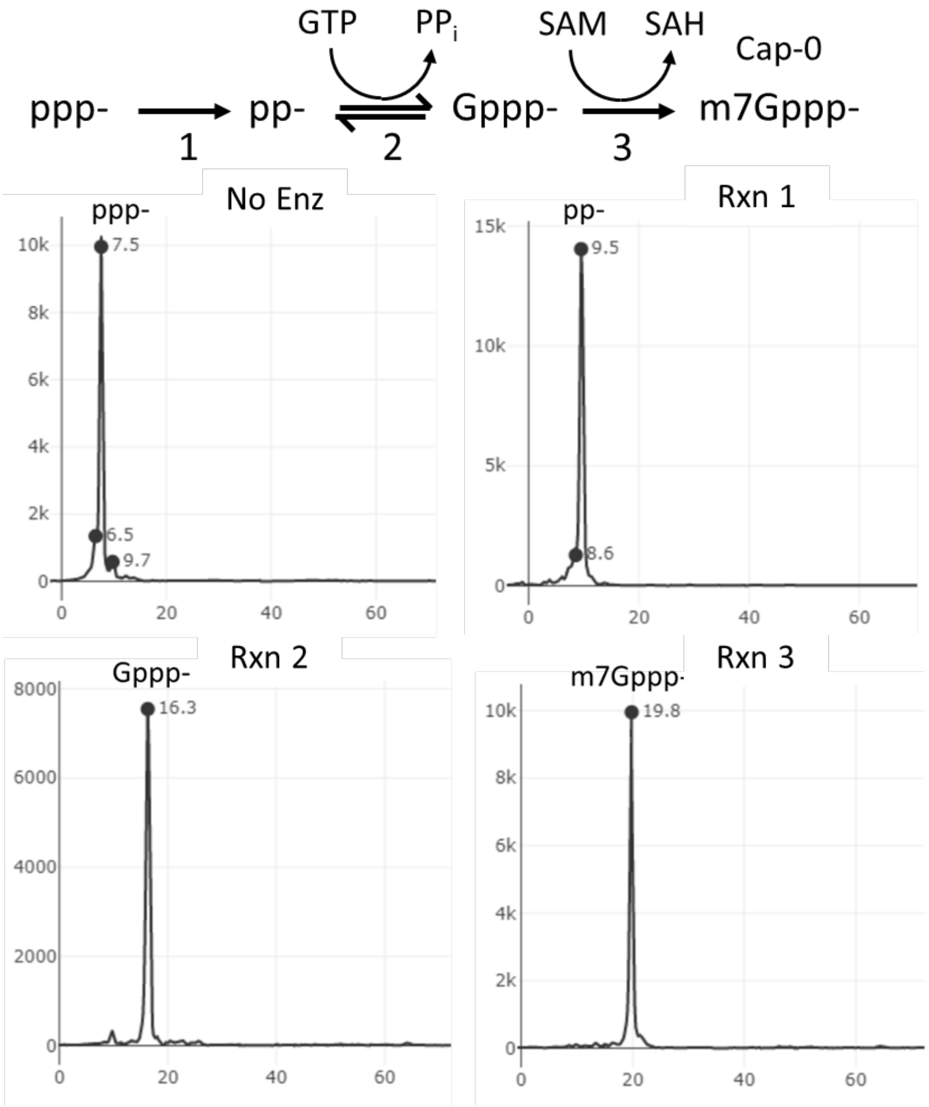


| Rxn | ppp25mer[FAM] | FCE | GTP | SAM | Expected product |
|---|---|---|---|---|---|
| No enz | + | - | - | - | ppp- |
| 1 | + | + | - | - | pp- |
| 2 | + | + | + | - | Gppp- |
| 3 | + | + | + | + | m7Gppp- |
